# Translational control of *gurken* mRNA in *Drosophila* development

**DOI:** 10.1101/076091

**Authors:** Christopher J. Derrick, Timothy T. Weil

**Affiliations:** Department of Zoology, University of Cambridge, Downing St, Cambridge, CB2 3EJ UK

## Abstract

Localised mRNA translation is a widespread mechanism for targeting protein synthesis, important for cell fate, motility and pathogenesis. In *Drosophila,* the spatiotemporal control of *gurken/TGF-α* mRNA translation is required for establishing the embryonic body axes. A number of recent studies have highlighted key aspects of the mechanism of *gurken* mRNA translational control at the dorsoanterior corner of the mid-stage oocyte. Orb/CPEB and Wispy/GLD-2 are required for polyadenylation of *gurken* mRNA, but mis-localised *gurken* mRNA in the oocyte is not fully polyadenylated^1^. At the dorsoanterior corner, Orb and *gurken* mRNA have been shown to be enriched at the edge of Processing bodies, where translation occurs^2^. Over-expression of Orb in the adjacent nurse cells, where *gurken* mRNA is transcribed, is sufficient to cause mis-expression of Gurken protein^3^. In *orb* mutant egg chambers, reducing the activity of CK2, a Serine/Threonine protein kinase, enhances polarity defects, consistent with a phenotype relating to a mutation of a factor involved in *gurken* translation^4^. Here we show that sites phosphorylated by CK2 overlap with active Orb and with Gurken protein expression. We also consolidate the current literature into a working model for *gurken* mRNA translational control and review the role of kinases, cell cycle factors and polyadenylation machinery highlighting a multitude of conserved factors and mechanisms in the *Drosophila* egg chamber.

## INTRODUCTION

The ability to regulate localised mRNA translation allows a cell exquisite spatiotemporal control over protein synthesis. The localisation, translation and degradation of mRNA requires the recruitment of trans-acting factors to the transcript, most commonly achieved through proteins binding to the 5′ and 3′ untranslated regions (UTRs). Once localised, initiation of translation involves scaffold factors and ribosomes being recruited to the mRNA by the 5′ cap and a sufficiently long 3′ poly(A) tail^5^. Multiple mechanisms exist that can regulate when and where the recruitment of these factors, required for translation takes place^6^.

One of the most common mechanisms is the packaging of transcripts into ribonucleoprotein complexes (RNPs) which contain RNA-binding proteins that prevent the recruitment of the initiation machinery^6^. RNPs serve to localise the mRNA through connecting with the cytoskeleton and after localisation is completed, translation is usually enabled through de-repression^6^ as exemplified by *ASH1* mRNA in budding yeast and *β-actin* mRNA in chicken fibroblasts.

During division of the budding yeast *Saccharomyces cerevisiae, ASH1* mRNA is localised to the tip of the future daughter cell where the protein acts to repress mating type switching^7^. Translation of *ASH1* mRNA is repressed in the mother cell through the binding of Puf6, inhibiting eIF5B function, thus blocking ribosome recruitment^8^. In the daughter cell, *ASH1* mRNA is de-repressed through CK2-dependent phosphorylation of Puf6, leading to localised translation^8^.

A similar mechanism functions at the leading edge of cultured migrating chick fibroblasts. Within the 3′ UTR of *β-actin* mRNA, the ‘zip-code’ sequence is bound by Zip-code binding protein 1 (ZBP1), which acts to both localise and repress the translation of the mRNA^9^. Once at the leading edge, the spatially restricted activity of Src kinase results in a change of ZBP1 binding affinity, allowing for localised *β-actin* translation^10^.

In the *Drosophila* egg chamber, localised translation of *gurken/TGF-α (grk)* mRNA is required for setting up the future body axes^11–13^. During mid-oogenesis (stages 7-10a), *grk* mRNA is transcribed and packaged into RNPs in the nurse cells, support cells that are interconnected with each other and the developing oocyte^14^. These *grk* mRNA containing RNPs are actively transported by the molecular motor dynein on microtubules to the dorsoanterior corner^15,16^, where translation and subsequent secretion of Grk protein patterns the dorsoventral axis^13,17^.

The translational repression of *grk* mRNA in the nurse cells was recently shown by genetic analysis to not require any previously described repressors^3^. Instead, the absence of the translational activator oo18-RNA binding protein (Orb), the *Drosophila* homolog of cytoplasmic polyadenylation element binding protein (CPEB) in the nurse cells prevents *grk* translation^3^. Although there is no direct evidence of Orb binding *grk* mRNA, examination of the 3′UTR shows that there are six weakly conserved cytoplasmic polyadenylation elements (CPEs) and in weak *orb* allele egg chambers *grk* mRNA is not fully polyadenylated^1^. Orb protein is restricted to the oocyte by an auto-regulatory loop dependent on the activity of Cup, an eIF4E binding protein (4E-BP) and dFMR1 (the *Drosophila* homolog of Fragile X Mental Retardation Protein) in the nurse cells^18–22^. This absence of an activator supports a model where in the nurse cells, *grk* mRNA is synthesised then exported to the oocyte with a short poly(A) tail maintaining translational repression until the mRNA is localised at the dorsoanterior corner. Sufficient levels of active Orb at the dorsoanterior corner are able to direct cytoplasmic polyadenylation, resulting in localised activation of *grk* translation ^1,3^. This polyadenylation is thought to be achieved by the *Drosophila* GLD-2 homolog Wispy, as the loss of cytoplasmic polyadenylation in *orb* mutant egg chambers is enhanced further by mutations in *wispy*^1^.

Egg chamber polarity, generated by the localised translation of *grk* mRNA at the posterior pole in stage 6 oocytes is required for the localisation of other axis-patterning maternal mRNAs^23^. One such mRNA is *bicoid (bcd),* the anterior determinant, which is localised along the anterior margin of the oocyte, partially overlapping with *grk* mRNA^24,25^. However, *bcd* mRNA remains translationally silent during oogenesis. This differential control of translation is regulated through association with Processing bodies (P bodies)^2^, electron dense, non-membrane bound structures, conserved in eukaryotes, that do not support translation through lack of ribosomes.

At dorsoanterior corner, where both *bcd* and *grk* mRNA are localised, the core of P bodies are enriched in translational repressors, whilst the periphery or edge (70nm in width) is enriched for the translational activator Orb^2^. *bcd* mRNA is concentrated six-fold at the P body core, whereas at the outer edge, *grk* mRNA is 23-times more concentrated^2^. This leads to a model whereby the edge of the P body (where *grk* mRNA is localised) is able to support translation through access to Orb and ribosomes, whilst the core (where *bcd* mRNA is localised) is not^2^. Compared to the oocyte, nurse cell P bodies do not associate with *grk* mRNA and have significantly reduced levels of Orb^3^. Over-expression of Orb in the nurse cells results in *grk* mRNA and Orb associating with the edge of P bodies and mis-expression of Grk protein in the nurse cells^3^. This result further supports that it is the restricted access to Orb that maintains *grk* mRNA repression in the nurse cells^3^.

Whilst access to Orb explains the lack of translational activation in the nurse cells, within the oocyte, Orb protein is not localised solely at the dorsoanterior corner^3^. Added to this, that mis-localised *grk* mRNA is not polyadenylated, it suggests that there is a mechanism regulating the activity of Orb^1^. One potential mechanism could be through localising the activity of Orb to the dorsoanterior corner through phosphorylation, a conserved way of regulating CPEB function^26–28^. Supporting this model is the presence of multiple phospho-isoforms of Orb in the egg chamber, which are altered or lost in mutants for either the catalytic or regulatory subunit of the Ser/Thr kinase CK2^4^. In wild-type oocytes, hyper-phosphorylated Orb associates with Wispy, an homologous interaction that occurs in the oocytes of *Xenopus laevis* ^4^,^28^,^29^. These observations, together with data showing that partially reducing CK2 activity in *orb* egg chambers enhances dorsoventral polarity defects^4^, suggests that CK2 is phosphorylating Orb, and this is required for *grk* translation.

Here we present new data to suggest that the localised translation of *grk* mRNA at the dorsoanterior corner is achieved through the spatial restriction of CK2 activity. This leads to phosphorylated Orb associating with *grk* mRNA at the P body edge where it directs cytoplasmic polyadenylation leading to Grk protein synthesis. We take this opportunity to also review the literature and present a working model for *grk* mRNA translation, with special consideration for components conserved with other systems.

## RESULTS

### CK2 activity is restricted to the oocyte and co-localises with active Orb

CK2 is a highly promiscuous enzyme, with a consensus site that has been hard to define. Various studies have shown that phosphorylation by CK2 is dictated by multiple acidic residues mostly downstream of the acceptor amino acid where n+3 is the most crucial^30^. A minimum consensus of S/T-X-X-E/D has been suggested, with the second most important residue (n+1), also normally acidic^30^. To investigate whether CK2 phosphorylates Orb, an antibody which recognises sites phosphorylated by CK2: pS/pT-D-X-E (where p represents a phosphorylated residue) was used in combination with an Orb antibody (see Methods).

At stage 6, when *grk* is localised at the posterior of the oocyte, Orb is present in the same pattern as *grk* mRNA and CK2 phosphorylation sites (Figure 1A-A″). Grk, secreted at the posterior of the oocyte binds Torpedo (Top) receptors on the overlying follicle cells. This Grk-Top signalling event polarises the egg chamber^23,31^, and leads to the oocyte nucleus being pushed to the future dorsoanterior corner^32^. *grk* mRNA becomes re-localised in a characteristic crescent at the dorsoanterior corner and then translated^11^. At stage 8, whilst Orb is present throughout the oocyte (Figure 1B) CK2 activity appears to be largely restricted to the dorsoanterior corner (Figure 1B″), where *grk* mRNA is known to be translated^12^. During both *grk* mRNA translation events, CK2 activity to be restricted only to the oocyte, with very little signal in the nurse cells (Figure 1A′ and B′). Together with genetic data where *ck2* mutant alleles in combination with *orb* mutants lead to dorsoventral defects in the oocyte^4^, our data strongly suggests that CK2 is acting to phosphorylate Orb at the dorsoanterior corner. We propose that the phosphorylated, active Orb at the dorsoanterior corner of the mid-stage oocyte is analogous to the previously described hyper-phosphorylated Orb^4^.

**Figure 1.**
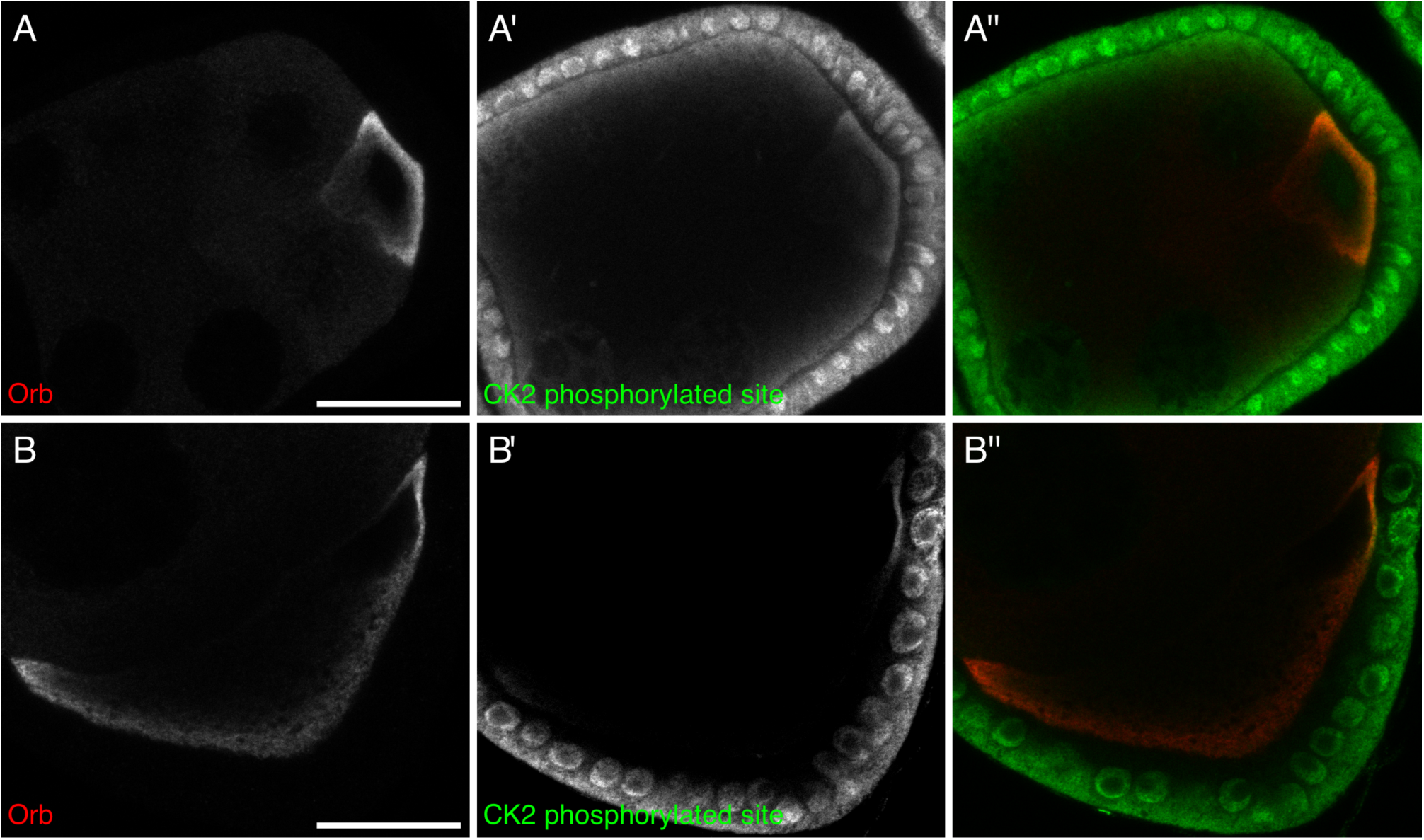
CK2 phosphorylation is restricted to the oocyte and overlaps with active Orb. Double immunofluorescent staining of egg chambers for Orb protein (red) and phosphorylated CK2 consensus sites (pS/pT-D-X-E, green) at stage 6 (A-A″) and stage 8 (B-B″) of oogenesis. **(A), (B)** Orb protein is localised throughout the developing oocyte. **(A′)** In stage 6 oocytes, sites phosphorylated by CK2 are present mainly at the posterior of the oocyte, between the nucleus and membrane, where localised *grk* translation is known to occur. There is minimal signal in the nurse cells, but all follicle cells show intense staining (n=20/20). **(B′)** In stage 8 oocytes, sites phosphorylated by CK2 are present almost exclusively at the dorsal-anterior corner, where *grk* mRNA is locally translated. Again there is little or no signal in the nurse cells and all follicle cells show intense staining (n=20/20). **(A″), (B″)** Merge of two staining showing that CK2 phosphorylated sites overlap with Orb protein at sites of localised *grk* translation, whilst regions of the oocyte that do not support *grk* translation are devoid of CK2 phosphorylated Orb. Single slices, scale bar 20pm.

### CK2 activity co-localises with Grk protein

The functional CK2 enzyme is a hetero-tetramer composed of two catalytic subunits (α) and two regulatory subunits (β)^33^. When either of these subunits are mutated to eliminate function (even when heterozygous) there is a change in the phosphorylation state of Orb^4^. There is also an increase in the percentage of dorsalised chorions in mature oocytes^4^, characteristic of a *grk* mutant phenotype.

Having shown that CK2 activity overlaps active Orb at the dorsoanterior corner (Figure 1B-B″), it would be expected that if CK2-dependent phosphorylation of Orb is directing localised *grk* translation, Grk protein would also overlap with CK2 phosphorylation. Double staining for CK2 phosphorylated targets and Grk protein in stage 8 chambers, shows that secreted Grk protein overlaps with the localised activity of CK2 (Figure 2), further implicating a role for CK2 in directing localised *grk* translation.

**Figure 2.**
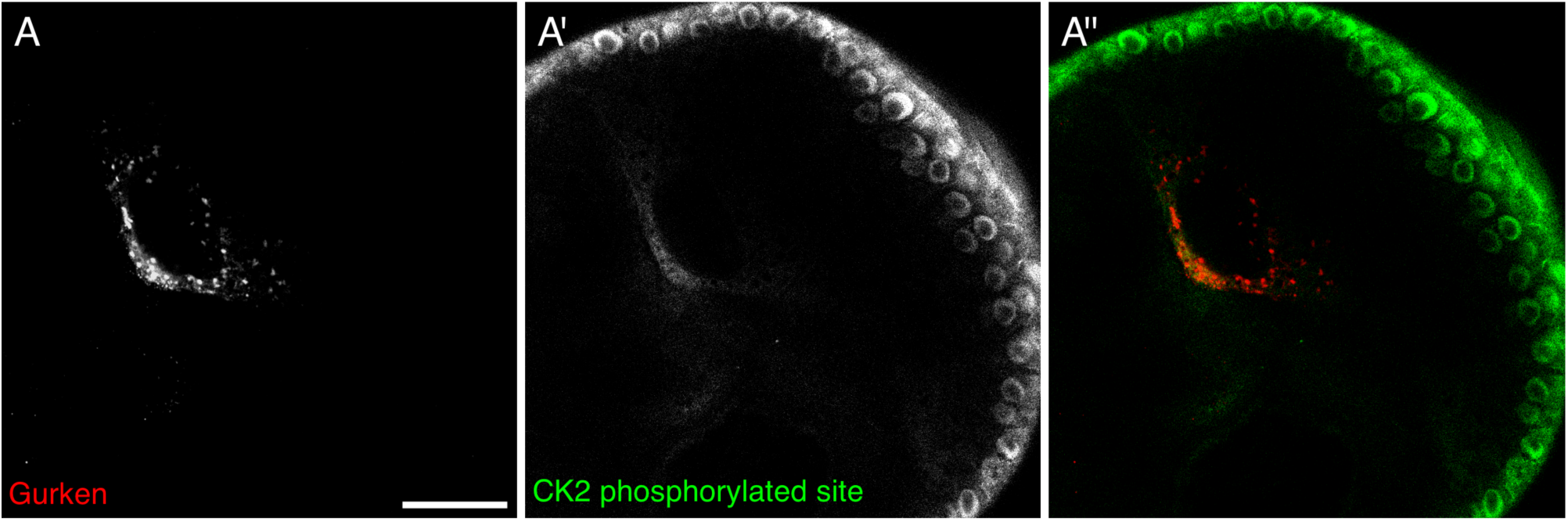
CK2 phosphorylation overlaps with Grk protein.

Double immunofluorescent staining of egg chambers for Grk protein (red) and phosphorylated CK2 consensus sites (pS/pT-D-X-E, green) at stage 8 of oogenesis. **(A)** Grk protein is present at the dorsal-anterior corner of stage 8 oocytes (n=15/15). **(A ′)** CK2 phosphorylated sites are present at the dorsal-anterior corner and in the surrounding follicle cells, but absent from nurse cell cytoplasm and other regions of the oocyte (n=15/15). **(A″)** Merge of two stains shown that phosphorylated CK2 consensus sites overlap with Grk protein secretion (n=15/15). Single slices, scale bar 20pm.

### CK2 activity co-localises with ectopically expressed Orb in nurse cells

The access to Orb in the nurse cells, adjacent to the developing oocyte is restricted through the function of the *orb* auto-regulatory loop which confines the majority of Orb protein to the oocyte through the action of Cup and dFMR1 in the nurse cells^18,19,22^. This repression is thought to be saturated when *orb* is over-expressed in the nurse cells since UAS-Orb egg chambers have ectopic Orb present in the nurse cells in a similar manner to that of *cup* mutants^3^. The over-expressed Orb is present in puncta at the edge of P bodies, along with *grk* mRNA which undergoes ectopic translation^3^. If CK2 activity is required to cause localised activation of Orb at the dorsoanterior corner in wild-type egg chambers, we hypothesised that in UAS-Orb egg chambers, the over-expressed Orb in the nurse cells, is phosphorylated by CK2.

Orb over-expression, as previously described^3^, results in Orb present in nurse cells in large foci and at a significantly higher level compared to wild-type egg chambers (compare Figure 3A to Figure 3B)^3^. We find that CK2 phosphorylation is present in the nurse cells of UAS-Orb egg chambers, whereas in wild-type nurse cells, CK2 activity does not appear to be present (compare Figure 3B′ to Figure 3A′ Moreover, the CK2 phosphorylation is detected overlapping with large foci of Orb (Figure 3B″, white arrows). Together these data strongly suggest a model whereby *grk* mRNA translation is directed through the localised activity of Orb at the edge of P bodies, achieved by localised CK2 activity.

**Figure 3.**
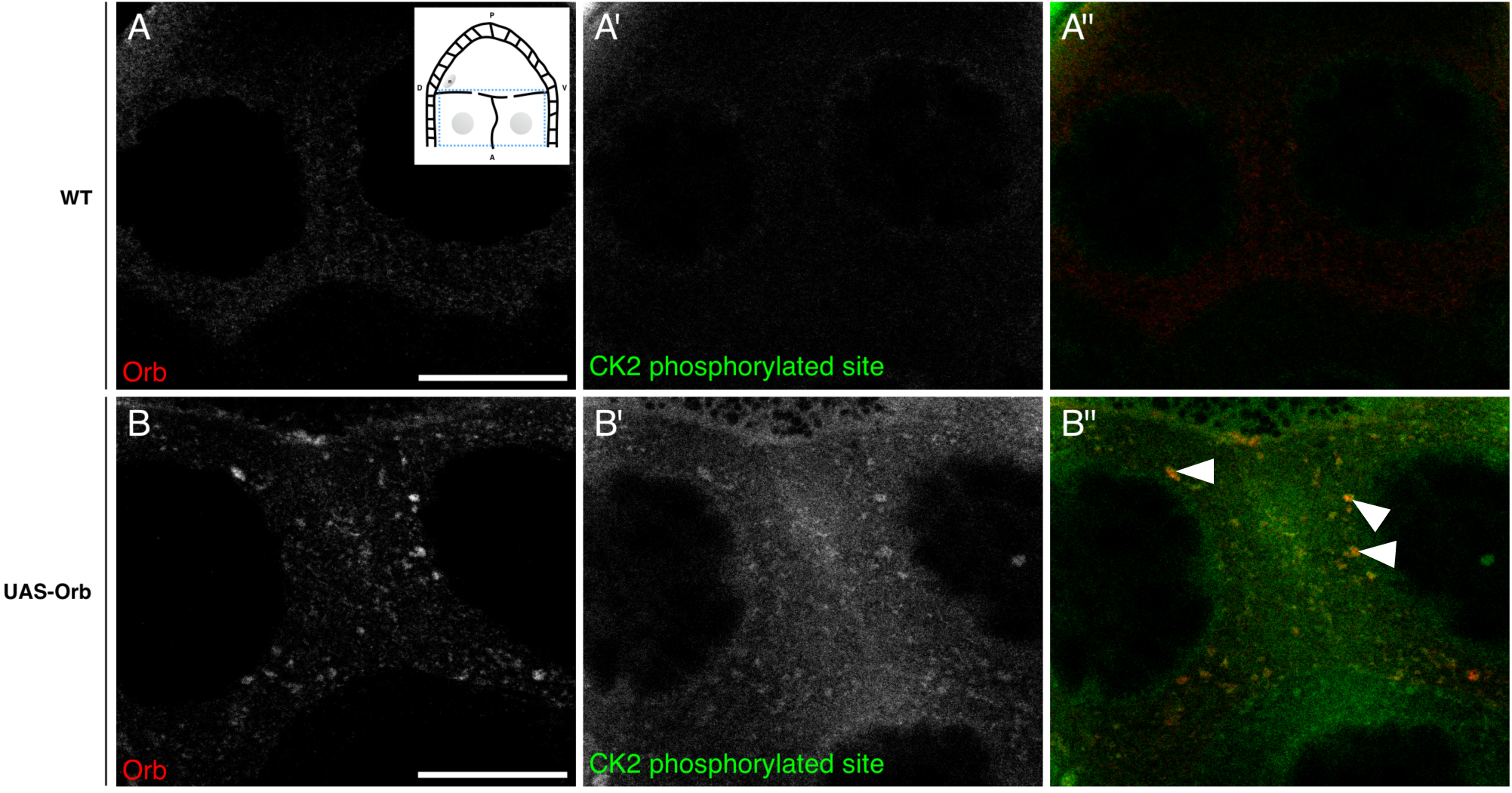
In UAS-Orb chamber, ectopically expressed Orb is phosphorylated by CK2. Double immunofluorescent staining of wild type (A-A″) and tubGal4-VP16; UAS-Orb (B-B″) egg chambers for Orb protein (red) and phosphorylated CK2 consensus sites (pS/pT-D-X-E, green). Inset: schematic illustrating the relative position of the oocyte and nurse cells for a stage 8 egg chamber, blue box highlights area focussed on in both (A-A″) and (B-B^″^). **(A-A″)** In the nurse cells of wild-type egg chambers, Orb protein is at a low level and CK2 activity is below the level of detection (n=12/12). **(B)** tubGal4-VP16; UAS-Orb egg chambers, have ectopically expressed Orb in foci or varying sizes throughout the nurse cells (n=12/12). **(B′)** Phosphorylated CK2 consensus sites are visible in tubGal4-VP16; UAS-Orb egg chambers (n=12/12). **(B″)** Merge of two stains showing that ectopically expressed Orb protein in the nurse cells is phosphorylated by CK2 (white arrowheads). Single slices, scale bar 20μm.

### Working model for *grk* translational control

Together with previous biochemical data and genetic interaction studies^4^, our data demonstrates that CK2 is phosphorylating Orb protein at the dorsoanterior corner during mid-oogenesis. Phosphorylation acts to direct Orb to the edge of P bodies, where *grk* mRNA is localised, resulting in translation. Coupled with previous work in this field, we suggest an updated model for the translational regulation of *grk* mRNA (Figure 4).

**Figure 4.**
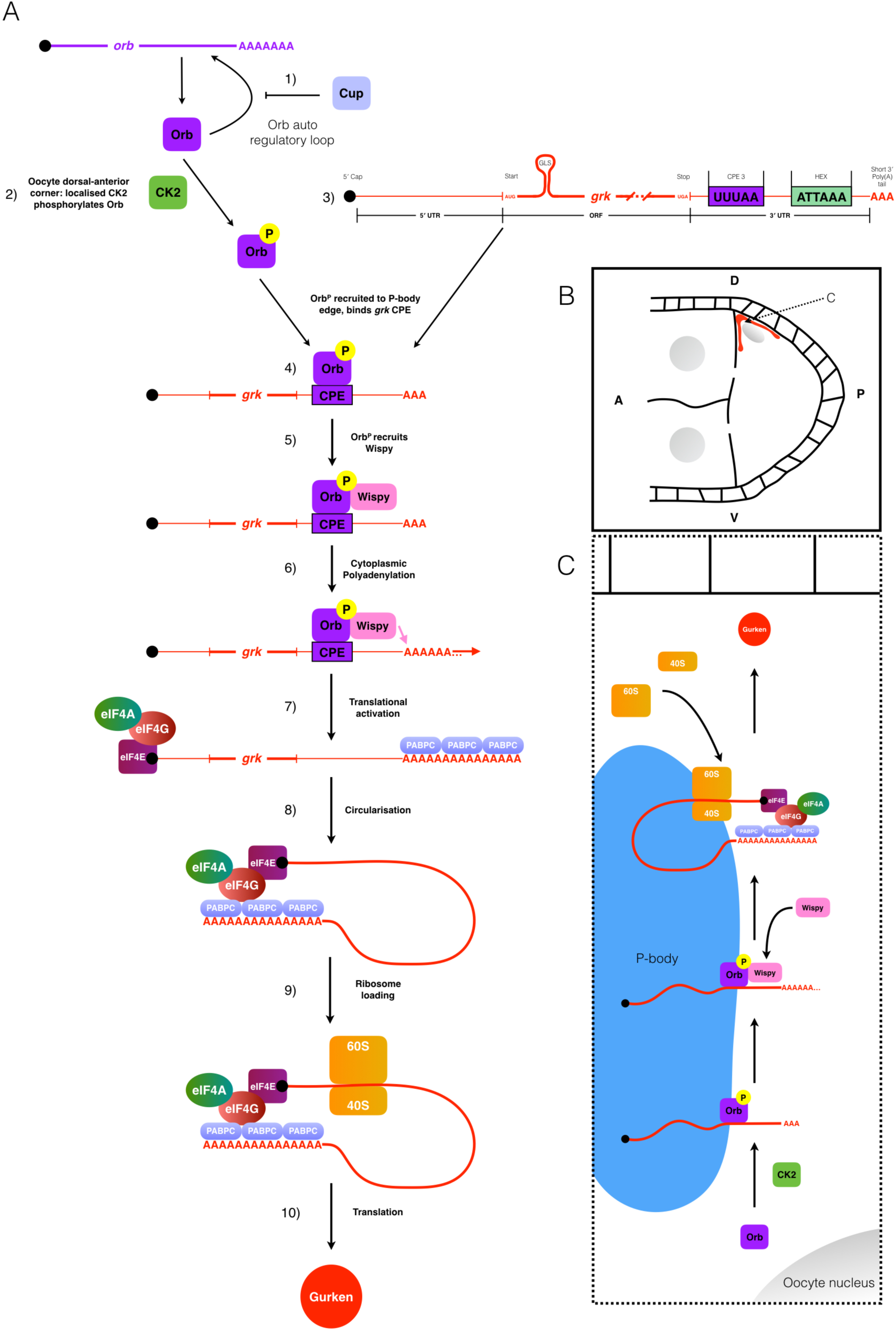
Model for *grk* translational control. Schematic collating data from this and other studies (references refer to the text) to present a model for the translational control of *grk* mRNA during stage 8 of oogenesis. **(A)** 1) *orb* mRNA translation is repressed in the nurse cells though the action of Cup^3,19^. 2) Orb is present throughout the oocyte, but is locally phosphorylated by CK2 at the dorsal-anterior corner. 3) Also present at the dorsal-anterior corner is *grk* mRNA, localised by the *gurken* localisation signal (GLS). 4) Phosphorylated Orb binds *grk* mRNA, presumably at one or many of the 6 weak CPE consensus sites present in the mRNAs 3′ UTR (only the strongest CPE is shown: CPE3)^1^. 5) Phosphorylated Orb is able to recruit the *Drosophila* homolog of the GLD-2 cytoplasmic polymerase Wispy^4,9^. 6) Wispy catalyses the addition of around 60As to the 3′ tail of *grk* mRNA extending it's length from 30 to 90As^1^. 7) Cytoplasmic polyadenylation of *grk* mRNA results in translational activation: canonical translation initiation machinery binds the mRNA 8) The mRNA is circularised through interactions between eIF4G and PABPC for efficient translation 9) The ribosome is loaded onto the mRNA. 10) Translation of *grk* mRNA generates Grk protein. **(B)** Stage 8 egg chamber with Anterior (A) left and Dorsal (D) up. Grk protein (red) is synthesised only at the dorsal-anterior corner, in a characteristic crescent between the oocyte nuclear (grey) and membrane. **(C)** Model for the mechanism of *grk* translational control at the dorsal-anterior corner. *grk* translational activation occurs at the edge of P-bodies^2^ at the dorsal-anterior corner of the oocyte. Phosphorylation of Orb by CK2 directs the CPEB homolog to the edge of P-bodies where *grk* mRNA is localised^2^. Orb then directs cytoplasmic polyadenylation resulting in translational activation of *grk* mRNA (see (A) for mechanistic detail).

In nurse cells the levels of the Orb protein are very low due to the action of the *orb* auto-regulatory loop (1, Figure 4A)^18,19^. This lack of Orb prevents *grk* mRNA translation in the nurse cells^3^. In the oocyte, Orb protein is present at a significantly higher level (2, Figure 4A)^3^. Orb is only targeted to the edge of P bodies at the dorsoanterior corner due to localised CK2-mediated phosphorylation (2, Figure 4A & Figure 4C)^2,3^. Phosphorylated Orb binds to localised *grk* mRNA, also present at the P body edge (3, 4, Figure 4A), likely through one or more of the six weak CPE consensus sites in the 3′ UTR (3, 4, Figure 4A & 4C, only the strongest consensus site, CPE3 is shown)^1–3^. Phosphorylated Orb interacts with Wispy^^1^,4,29^ (5, Figure 4A & Figure 4C) which polyadenylates *grk* mRNA (6, Figure 4A & Figure 4C)^1^. The *grk* mRNA poly(A) tail increases in length from 30As to 90As^1^ (7, Figure 4A). The sufficiently long poly(A) tail of *grk* mRNA is then bound by the canonical translation initiation machinery leading to ribosome recruitment (8, 9, Figure 4A) thus resulting in localised Grk protein production at the dorsoanterior corner (10, Figure 4A, Figure 4B and 4C).

## DISCUSSION

Our work, and that of others, continues to demonstrate the powerful nature of the developing egg chamber as a model for translational control through highlighting conserved mechanisms.

### CPE-mediated translational control

Here we have shown similarities to other systems that regulate CPEB homologs to direct the cytoplasmic polyadenylation of specific mRNAs. Cytoplasmic polyadenylation allows for both spatial and temporal control of protein synthesis, separating translation from nuclear transport.

This method of translational control is critical in oocytes as these specialised cells are biologically “paused” with no transcription or translation for extended periods of time, in extreme examples, years. The silenced state of many maternal mRNAs with a short poly(A) tail is reversed by cytoplasmic polyadenylation at one of two major developmental events: oocyte maturation or the oocyte-to-embryo transition.

Oocyte maturation occurs following the main phase of growth and differentiation of the oocyte. Maturation allows for the co-ordination of the completion of meiosis with fertilisation. During this event, the cell cycle machinery is re-activated by the translation of a subset of mRNAs. This results in an active CDK-CyclinB complex which initiates and maintains a second pause in the cell cycle at metaphase until egg activation^34^.

Typically coupled with fertilisation, egg activation releases the egg from the pause in metaphase established at maturation and is associated with a transient rise in the concentration of intracellular calcium (Ca^2^+) levels^35^. Egg activation is part of the oocyte-to-embryo transition, when the mature, quiescent oocyte receives the necessary cues to form the totipotent, rapidly dividing early embryo.

Whilst the role for cytoplasmic polyadenylation was first described at egg activation in sea urchin eggs^36^, the majority of research on how cytoplasmic polyadenylation is regulated has been undertaken in *Xenopus* oocytes. This work has focused on the steps required to re-activate the translation of two different subsets of mRNAs at oocyte maturation, the passage from meiotic prophase I to metaphase II.

RNA injection experiments in maturing *Xenopus* oocytes revealed a *cis-element* within mRNAs known to undergo cytoplasmic polyadenylation, termed the cytoplasmic polyadenylation element (CPE). The CPE is U-rich and roughly octameric, with a consensus of U_4-5_A_1-3_U^37,38^ To determine the potential CPE binding protein, radio-labelled CPE-containing mRNA was incubated in *Xenopus* extracts and then irradiated to cross-link bound proteins for purification^38^. This identified cytoplasmic polyadenylation element binding protein (CPEB) an RNA-binding protein essential for the cytoplasmic polyadenylation of CPE containing mRNAs^38^. In vertebrates there are four CPEB proteins that share the conserved two RNA Recognition Motifs followed by a zinc finger domain in the C-terminus. For CPEB1, the originally isolated isoform, all three of these domains are required for RNA binding^38^. Whilst the N-termini are widely divergent between species, CPEB is evolutionary conserved, and has been identified in the worm, fly, fish, mouse, frog, a clam, a sea slug and human^39^.

The regulation of CPEB binding to CPE and subsequent recruitment of the poly(A) machinery must be regulatedsince CPEB is present in the cytosol along with CPE-containing mRNAs. This is most commonly achieved through post-translational modification, a wide-spread mechanism in post-transcriptional control.

*Xenopus* CPEB1 has been shown to undergo a number of phosphorylation events during maturation. Progesterone signalling from the surrounding follicle at maturation activates Eg2 (related to Aurora A Kinase), part of the events to co-ordinate further oocyte development^40,41^. Phosphorylation by Eg2 strengthens the interaction between CPEB1 and xGLD-2 (the *Xenopus* homolog of the cytoplasmic poly(A) polymerase)^28^, leading to poly(A) tail extension and subsequent translation. Later during maturation, a second phosphorylation event, which is dependent on CDK-CyclinB, occurs at S210 of CPEB1^42^. This phosphorylation targets most, but not all, of CPEB1 for proteolytic degradation and drastically reduces the ratio of CPEB1 to CPEs in the cytoplasm. This is necessary to allow a second subset of mRNAs to undergo translational activation in the final steps of oocyte maturation^43–45^.

*Drosophila* oocyte maturation reveals some conservation of mechanism with *Xenopus.* Cortex (Cort) is a distant member of the Cdc20 family of APC/C (Anaphase Promoting Complex/ Cyclosome) specificity factors and is required for female meiosis^46,47^. Cort functions through binding APC/C (forming APC/C^Cort^) to direct the proteolytic activity ensuring that the cell cycle progresses correctly^48^. Cort is not detectable during early oogenesis, but is present at the beginning of maturation (stage 13 of oogenesis) and remains active until egg activation^34,49^. The appearance of Cort and the activity of APC/C^Cort^ at maturation coincides with both the cytoplasmic polyadenylation of *cort* mRNA and the degradation of CyclinA, which is necessary for entry into metaphase^49,50^.

The cytoplasmic polyadenylation of *cyclinB (cycB)* mRNA occurs concurrently with *cort* mRNA^49^. APC/C^Cort^ at maturation importantly only targets CyclinA. At this time, CycB and Securin are not targeted and can maintain the mature *Drosophila* oocyte in metaphase I^49^. For both *cycB* mRNA and *cort* mRNA the cytoplasmic polyadenylation is dependent on Wispy^29^. Altogether, this demonstrates the high level of conservation involved in cytoplasmic polyadenylation at oocyte maturation for *Drosophila* and *Xenopus.* However, it has not been shown if *Drosophila cort* and *cycB* mRNA possess CPEs.

Cytoplasmic polyadenylation is not limited to the germ-line. In mouse neurons, Calmodulin-dependent Kinase II (CaMKII) dependent phosphorylation of CPEB at T171 (a site highly similar to *Xenopus* CPEB1 S174) is able to regulate translational activation^26^. This co-ordination of translation of CPE containing mRNAs in response to increased intracellular calcium facilitates synaptic plasticity^26^. *CaMKII* mRNA is also known to undergo cytoplasmic polyadenylation mediated by Aurora kinase-catalysed CPEB phosphorylation at synapses in response to external cues^27^.

CPEB homologs have also been implicated in disease through their role in directing localised translation. CPEB1 in mammary glands is required to correctly localise mRNAs encoding polarity proteins that are required for the construction of tight junctions. CPEB1 absence in these cells results in the loss of apico-basal polarity^51^, which can lead to the epithelial-to-mesenchymal transition (EMT) resulting in metastasis^52^. Depletion of CPEB1 can also lead to cancer progression by directly altering poly(A) tail lengths of EMT/metastasis-related mRNAs^53^. One example, *Matrix metalloproteinase 9* (*MMP9*) mRNA, encodes a metastasis-promoting factor^54^ and in CPEB1 depleted cells, there is an increase in the poly(A) tail length of *MMP9* that results in translational activation^53^.

### *grk* mRNA as a model for CPE-mediated translational control

The regulation of Orb in *grk* translational control appears to be very distinct from that of CPEB1 regulation in *Xenopus* oocytes. In *Xenopus,* CPEB1 is believed to be weakly associated with mRNA prior to maturation and Eg2 phosphorylation strengthens this and other interactions, with factors required for cytoplasmic polyadenylation such as xGLD-2^28^. However, we have demonstrated that phosphorylation of the CPEB homolog Orb at the dorsoanterior corner is important to localise Orb with *grk* mRNA to direct cytoplasmic polyadenylation at the edge of P bodies. Therefore for *grk* translational control, phosphorylation of Orb appears to regulate the activity of the protein spatially, rather than temporally. This reflects a key difference in the mRNAs undergoing translational control in these systems, Grk protein must be secreted in the correct place for patterning, however for mRNAs in maturing *Xenopus* oocytes, the polyadenylation must occur at the right time to ensure cell cycle reactivation.

This localised translation of *grk* mRNA does resemble the translational control of *ASH1* mRNA in budding yeast and *β-actin* mRNA in migrating chick fibroblasts. In both these examples, the activity of a kinase is localised and required to initiate the translation of these mRNAs in the correct sub-cellular region^8,10^. Contrastingly to mRNAs undergoing translational activation in the maturing *Xenopus* oocyte, their spatial control is much more important than their temporal control.

What is particularly interesting is the conserved use of localised CK2 activity between *Drosophila* and *Saccharomyces cerevisiae*, despite the two systems being separated by millions of years of evolution. CK2-GFP in yeast is present throughout the dividing cells, but localises strongly in the cortex of the future daughter cell, prior to the expression of *ASH1* mRNA^8^. It is likely that this mechanism is similar in the *Drosophila* oocyte, with CK2 localised at the dorsoanterior corner to direct Orb phosphorylation in the correct position to establish dorsoventral polarity.

However, this presents a further set of questions: how is CK2 activity localised first at the posterior and then relocalised at the dorsoanterior corner for dorsoventral axis establishment? How is CK2 activity restricted to the oocyte only, given that in UAS-Orb egg chambers CK2 phosphorylates the ectopically expressed Orb? It would seem most likely that CK2 is active in the whole egg chamber but due to the low levels of Orb in the nurse cells, the signal is not visible and ultimately no *grk* translation occurs. Thus the spatial restriction of Orb to the oocyte is key to regulating *grk* mRNA translational control.

### The role of the *orb* auto-regulatory loop in *grk* translational control

During the early stages of oogenesis, Orb is critical for the specification of the oocyte^55^. Orb preferentially accumulates in the oocyte due to an auto-regulatory loop where Orb is required to localise its own mRNA^18^. Concurrently factors act in the nurse cells to prevent the incorrect activation of the auto-regulatory loop. One key repressor is Cup, a 4E-BP^19^. In *cup* mutants the *orb* auto-regulatory loop is not confined to the oocyte, resulting in high levels of Orb protein in the nurse cells^3,19^. Our previous work has shown that this loss of Orb restriction solely to the oocyte is sufficient to result in the ectopic translational of *grk* mRNA in nurse cells that leading to dorsoventral defects^3^. This suggests an indirect role of Cup in the translational control of *grk* mRNA^3^ whilst previous work, prior to the full elucidation of the *orb* auto-regulatory loop suggested a direct role for Cup in binding *grk* mRNA^56^. Therefore, with our UAS-Orb data presented here, and based on previous work, a complete understanding of the *orb* auto-regulatory loop is important to understand how *grk* translational control is achieved in the developing oocyte.

In *cup* mutants, *orb* mRNAs have longer poly(A) tails and the majority of Orb protein is hyper-phosphorylated^19^. However the role of Cup as a 4E-BP does not seem important for regulating the translation of *orb,* as specific loss of the eIF4E binding domain in *cup^Δ212^*does not have a strong effect on dorsoventral polarity as would expected if this were the sole role of Cup (Davidson, unpublished). This would suggest that there are functions for Cup other than eIF4E sequestration that restrict the *orb* auto-regulatory loop to the oocyte. One potential mechanism by which Cup could be acting to repress *orb* translation in the nurse cells, fitting the elongated poly(A) tails, is through promoting the recruitment of the deadenylation machinery to the mRNA^57^.

In both *cup* and UAS-Orb egg chambers, Grk is expressed along the margins of nurse cells although less severely in UAS-Orb^3^. In UAS-Orb nurse cells, we have shown that Orb co-localises with CK2 phosphorylated sites (Figure 3) and previously shown that this ectopic Orb is present at the edge of nurse cell P bodies resulting in *grk* translation^3^. The phosphorylation of Orb by CK2 in the nurse cells when over-expressed, would fit with the increased amount of hyper-phosphorylated Orb in *cup* mutants^19^. Together this would suggest that Cup-mediated repression of *orb* translation in the nurse cells is not only directly important for oocyte specification, but also indirectly for the translational control of *grk* mRNA.

Loss of Orb repression does not fully explain the *cup* phenotype as in *cup*^5^, a particularly strong allele, *grk* mRNA is less efficiently localised^56^. Another component shown to be involved in the control of *grk* translation is the heterogeneous nuclear ribonucleoprotein (hnRNP) Squid (Sqd)^58–60^. Sqd and Cup have been shown to interact through both biochemical and genetical analysis where *cup* mutants enhance the dorsalised phenotype of *sqd* homozygotes^56^. However, whether this enhancement of *sqd* by *cup* is specifically due to loss of function of Cup independent of the role in the *orb* auto-regulatory loop remains to be fully investigated.

*sqd*^1^ mutants show Grk protein expressed along the entire anterior margin of the developing oocyte^61^. How specifically Sqd functions remains unclear, particularly as this loss of *sqd* does not lead to ectopic expression in the nurse cells^3^. Loss of *sqd* does results in a failure to localise *grk* mRNA to the dorsoanterior corner and leads to ectopic translation along the anterior margin^3,58,61^. This suggests that there is a mechanism involved in controlling the translational activation of *grk* mRNA in the oocyte that is dependent on Sqd. In light of the data on the role of Cup in the *orb* auto-regulatory loop, it seems that the roles of Sqd and Cup in *grk* translational control in the oocyte need to be re-considered.

### The role of Bruno in *grk* mRNA translational control

Further suggesting that there is not a role for a translational repressor of *grk* mRNA in the nurse cells comes from over-expression of the *grk* locus. In UAS-*grk* egg chambers, there is no ectopic *grk* translation in the nurse cells, suggesting that there is not a saturatable binding of repressors that occurs in the nurse cells to repress *grk* translation^3^. In these *grk* over-expression egg chambers, previous works shows that some excess *grk* mRNA is localised to the repressive P body core^2^. There is, however, some Grk over-expression as dorsoventral patterning defects, characteristic of a failure in *grk* translational control are observed^2,3^.

Previously a number of studies had implicated Bruno (Aret) in the translational repression of *grk* mRNA in the oocyte^56,61–63^. Over-expression of Bruno does not affect the localisation of *grk* mRNA, but does result in loss Grk protein^62^. Further evidence in support of Bruno-mediated repression showed that Bruno bound to the 3′ UTR of *grk* mRNA^62^. This is at odds with recent imaging data, where in different *aret* mutants, *grk* mRNA is not ectopically translated in the nurse cells or the oocyte^3^. Interestingly, Bruno has been shown to associate with presumed non-active hypo-phosphorylated Orb^4^, which we suggest is only present at low levels at the dorsoanterior corner, whilst previous work has shown Bruno is enriched at the dorsoanterior corner^64^. One potential explanation is that Bruno may function as a redundant repressor for the translational control of *grk* mRNA in the oocyte. Bruno would not be required in the nurse cells, due to the low levels of Orb. Complementing direct, *in vivo* visualisation and genetic approaches should begin to unravel these conflicting observations.

### Future Directions

Combining our work with others, we present a model of *grk* mRNA translational control that is dependent upon the localised activity of CK2 at the dorsoanterior corner of the mid-stage oocyte (Figure 4). The observation of the localisation of CK2 itself during the two periods of *grk* translation would further validate our own imaging and others genetic and biochemical data. Generation of a GFP-tagged CK2 would allow for visualisation of the location of the enzyme. This could also be used to follow how CK2::GFP is localised over the course of oogenesis to direct *grk* translation. Examining CK2::GFP localisation in nurse cells would also confirm our suggestion that CK2 is present and active in the nurse cells, but there is little Orb protein for the kinase to phosphorylate, due to the activity of Cup in the *orb* auto-regulatory loop.

The model also draws a number of similarities between the regulation of the poly(A) machinery in maturing *Xenopus* oocyte. In particular that the phosphorylation of Orb by CK2 recruits Wispy to *grk* mRNA at the edge of P bodies^4,29^. Direct visualisation of the interactions between Wispy and *grk* mRNA in wild-type and *ck2* backgrounds would begin to uncover the mechanism by which Wispy is recruited to *grk* mRNA.

This work continues to highlight the *Drosophila* egg chamber as a powerful model for elucidating mechanisms of mRNA translational control. The proposed model demonstrates that post-translational modification of CPEB homologs is a conserved mechanism, not only for temporal, but also spatial control of cytoplasmic polyadenylation.

## METHODS

### Fixed Oocyte Immunofluorescence

For all fixed oocyte immunofluorescence, the procedure was modified from Davidson et. al. 2016^3^ and imaged using a Leica SP5 inverted scanning confocal microscope equipped with Hybrid Detectors.

– Oocytes were dissected in Schneider's medium, ovarioles were splayed, but not separated
– Fixed for 10 minutes in 4% PFA (Alfa Aesar: J61899)
– Equal volume of Heptane was added for 3 minutes, vortexing intermittently
– 3 fast rinses followed by 3 slow washes in 0.2% PBST
– 1 wash in 1% PBS Triton-X 100 (Sigma X100)
– 1 rinse in 0.2% PBST
– Block in 30 minutes in 4% BSA (Sigma A7906)
– Primary antibodies incubation for 16hrs. at 4 degrees, overnight
– 3 fast rinses followed by 3 slow washes in 0.2% PBST
– Secondary antibodies added at room temperature for 1 hr.
– 3 fast rinses followed by 3 slow washes in 0.2% PBST
– DAPI
– Ovaries were mounted in Prolong Gold (Life technologies P36930), with any stages older than 9 removed and left at 4 degrees for a minimum of 16hrs. prior to imaging

| Target | Primary (Dilution) | Origin | Secondary (Dilution) |
| --- | --- | --- | --- |
| **Orb** | Mouse (1:40) | DSHB (Orb 4H8) | Donkey-647 (1:200) |
| **Grk** | Mouse (1:300) | DSHB (1D12-c) | Donkey-647 (1:200) |
| **Sites phosphorylated by CK2 (CK2 phospho)** | Rabbit (1:200) | CST (#8738) | Goat-Cy3 (1:200) |

Images were acquired using 100x, 0.4NA oil immersion lense with sequential settings

Sequential scan 1:

– DAPI: 405 Diode. Emission 405nm 3%. Collection 430-485nm. Gain: 130
– 647: He 633. Emission 633nm 4% (Orb) 15% (Grk). Collection 650-730nm. Gain: 115/130

Sequential scan 2:

– Cy3: DPS 561. Emission 561 nm 3%. Collection 570-610nm. Gain: 80

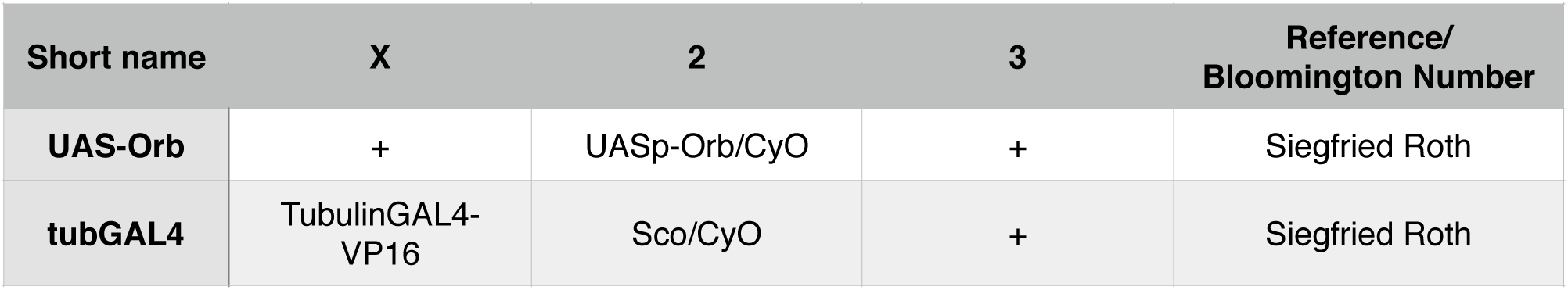

| Short name | X | 2 | 3 | Reference/<br>Bloomington Number |
| --- | --- | --- | --- | --- |
| **UAS-Orb** | + | UASp-Orb/CyO | + | Siegfried Roth |
| **tubGAL4** | TubulinGAL4-VP16 | Sco/CyO | + | Siegfried Roth |

